# Complex pleiotropic genetic architecture of evolved heat stress and oxidative stress resistance in the nematode *Caenorhabditis remanei*

**DOI:** 10.1101/515320

**Authors:** Christine H. O’Connor, Kristin L. Sikkink, Thomas C. Nelson, Janna L. Fierst, William A. Cresko, Patrick C. Phillips

## Abstract

The adaptation of complex organisms to changing environments has been a central question in evolutionary quantitative genetics since its inception. The structure of the genotype-phenotype maps is critical because pleiotropic effects can generate widespread correlated responses to selection and potentially restrict the extent of evolutionary change. In this study we use experimental evolution to dissect the genetic architecture of natural variation for acute heat stress and oxidative stress response in the nematode *Caenorhabiditis remanei*. Previous work in the classic model nematode *C. elegans* has found that abiotic stress response is controlled by a handful of genes of major effect and that mutations in any one of these genes can have widespread pleiotropic effects on multiple stress response traits. Here, we find that acute heat stress response and acute oxidative response in *C. remanei* are polygenic, complex traits, with hundreds of genomic regions responding to selection. In contrast to expectation from mutation studies, we find that evolved acute heat stress and acute oxidative stress response for the most part display independent genetic bases. This lack of correlation is reflected at the levels of phenotype, gene expression, and in the genomic response to selection. Thus, while these findings support the general view that rapid adaptation can be generated by changes at hundreds to thousands of sites in the genome, the architecture of segregating variation is likely to be strongly parcellated by the pleiotropic structure of the underlying genetic networks.

## Introduction

Biological organisms are complex, integrated systems. From an evolutionary point of view, a central consequence of this integration is that natural selection acting on one feature of an organism has the potential to have cascading effects across the whole organism (Lande 1979; Cheverud 1984; Phillips and McGuigan 2006). Evolutionary quantitative genetics has created a powerful statistical framework for analyzing complex phenotypes, even in the absence of knowledge of seemingly important factors such as the exact number or identity of genes or the distribution of their phenotypic effects (Barton and Turelli 1989). The advent of whole genome analyses would seem to herald the advent of a new era in which we have the power to understand the genetic basis of complex, multifactorial traits. However, even now, the field has achieved remarkably little success identifying the genetic basis of most adaptive phenotypes within natural populations, aside from traits where a majority of the variation is due to a handful of well-characterized mutations, such as coat color and oxygen affinity in deer mice (Steiner *et al*. 2007; Linnen *et al*. 2013; Natarajan *et al*. 2013). There may be a small handful of cases in which natural variation is generated by simple genetics, but it is clear that for many, if not most, traits this will not be the case (Phillips 2005; Rockman 2012). For example, human height is a well-studied, highly heritable, quantitative trait and yet despite sample sizes that now stretch into the millions, mapped loci still only account for a small fraction of the total genetic variance (Lango Allen *et al*. 2010; Nolte *et al*. 2017). Similarly, studies of complex diseases such as schizophrenia are able to map hundreds of genes associated with disease prevalence but still account for less than 10% of the variation in disease risk (Schizophrenia Working Group of the Psychiatric Genomics Consortium, 2014).

However, genes rarely affect only a single, isolated trait within an organism. The idea that genetic coupling across functional systems should be the rule rather than exception led Sewall Wright (1968) to posit the idea of “universal pleiotropy” for genetic systems. More recently, Boyle *et al*. (2017) proposed an ‘omnigenic’ model for complex diseases that posits that complex diseases could be affected by variation in genes in any tissue that affects that disease, i.e. such diseases are affected by variation in sets of core and peripheral genes. To a large extent, the omnigenetic paradigm is simply a translation of one of the central conceits of evolutionary quantitative genetics to the setting of human association mapping. At its limit, extensive pleiotropy can serve to “lock” the response to selection, creating a genetic constraint that limits the rate of evolutionary change (Lande 1979; Charlesworth 1990; Arnold 1992). On the other hand, variation in the functional specificity across the network of genes affecting complex traits, such as differences in tissue-specific regulation (Carroll 2008), holds the possibility that the genetic changes that actually lead to evolutionary divergence could be biased towards subsets of genes with either more or less pleiotropy, depending on the specific pattern of selection (Phillips and McGuigan 2006). Pleiotropy may therefore play a two-fold role in the response to selection. First, following traditional quantitative genetic approaches, patterns of pleiotropy underlying standing variation may lead to correlated responses to selection across broad suites of characters. Alternatively, following a developmental-genetic perspective, long-term evolutionary change may be strongly dependent on variation in pleiotropy, and may subsequently lead to a biased subset of genes exhibiting reduced pleiotropic effects, leading to the evolutionary change in of the pattern of pleiotropy itself (Phillips and McGuigan 2006; Arnold *et al*. 2008; Jones *et al*. 2014). The balance between these factors is largely an empirical question.

Several studies have found, depending on the organism, median levels of pleiotropy to be between 4–7 traits affected by every gene (Wagner and Zhang 2011; Paaby and Rockman 2013), but there are many cases in which a single mutation can affect dozens of traits (Knight *et al*. 2006). These studies tend to focus on what Paaby and Rockman (2013) have called “molecular gene pleiotropy”: how many functions does a given locus affect when that gene is knocked out. There is substantially less information on the extent of pleiotropy within populations generated by naturally segregating variation within populations, or “mutational pleiotropy” (Wagner and Zhang 2011; Paaby and Rockman 2013). Here the focus is on the entire spectrum of pleiotropy generated by segregating alleles as opposed to genetic knockouts. Thus it is the allele – and not the gene – that is the appropriate unit of variation for evolutionary change (Wright 1968; Lande 1984; Phillips and McGuigan 2006; Wagner and Zhang 2011). Studies that use forward genetic screens may find a large fraction of potential targets of those genes, but that does not necessarily inform us as to how a trait of interest will respond to selection.

How can we understand the balance between molecular and mutational pleiotropy, especially in the context of naturally segregating variation influencing complex correlated traits? Experimental evolution coupled with whole genome sequencing, or evolve and re-sequence, has emerged as a powerful method for studying the genetic architecture of complex traits (Kofler and Schlötterer 2013). Selection on complex traits can leave weak or ambiguous signals of selection on the genome (Kemper *et al*. 2014), but evolve and re-sequence facilitates direct comparison between ancestral and evolved populations and quantification of evolutionary change across the genome. In particular, replicated experiments within a well-defined hypothesis testing framework allow sampling variance to be accounted for in a way that historical contingency renders impossible within natural populations. While we have learned a great deal from experimental evolution in microbial populations initiated from a single fixed genetic background, questions related to the influence of complex segregating variation on the response to selection can only be addressed by capturing a broad array of natural genetic variation in the initial ancestor population (Schlotterer *et al*. 2015).

The nematode *Caenorhabditis remanei* is ideal for addressing questions relating to complex traits via experimental evolution (Sikkink, Reynolds, *et al*. 2014; Castillo *et al*. 2015; Teotónio *et al*. 2017). It is a genetically diverse, sexually reproducing organism (Jovelin *et al*. 2003; Cutter *et al*. 2006). Large populations can easily be maintained in a lab, with upwards of 2000 individuals living on one 10cm agar plate, so an entire experiment can occupy one shelf in an incubator. Additionally, its close relationship to the model nematode *C. elegans* allows both genomic and functional information to be applied across species.

Abiotic stress response in nematodes provides an interesting model system of interacting traits. In particular, many of the important stress response pathways are well conserved throughout animals and have important pleiotropic effects on aging and longevity (Kenyon 2010), with a handful of stress response pathways being know to affect the response to many different abiotic stressors (Rodriguez *et al*. 2013; Murphy and Hu 2013). For example, as elucidated within numerous studies in the well-studied insulin/insulin-like growth factor signaling pathway in *C. elegans*, mutations often affect resistance to a wide variety of stressors including heat stress, oxidative stress, ultraviolet radiation and pathogen stress (Murphy and Hu 2013). However, the role that natural genetic variation plays in individual variation in these traits is less clear (Reynolds and Phillips 2013; Jovelin *et al*. 2014).

Here, we use experimental evolution and whole genome sequencing to dissect the genetic architecture of acute heat stress and acute oxidative stress response in *C. remanei*. In previous work, we saw a strong phenotypic response to selection for both acute heat stress and acute oxidative stress but little to no correlated phenotypic response between selection for acute heat stress and acute oxidative stress resistance (Figure 1; Sikkink *et al*. 2015). In this study, we seek to align these global phenotypic patterns with a detailed analysis of gene expression and the genomic response to selection at single nucleotide resolution in order to test the complexity and coherence of genotypic and phenotypic responses to selection.

**Figure 1.**
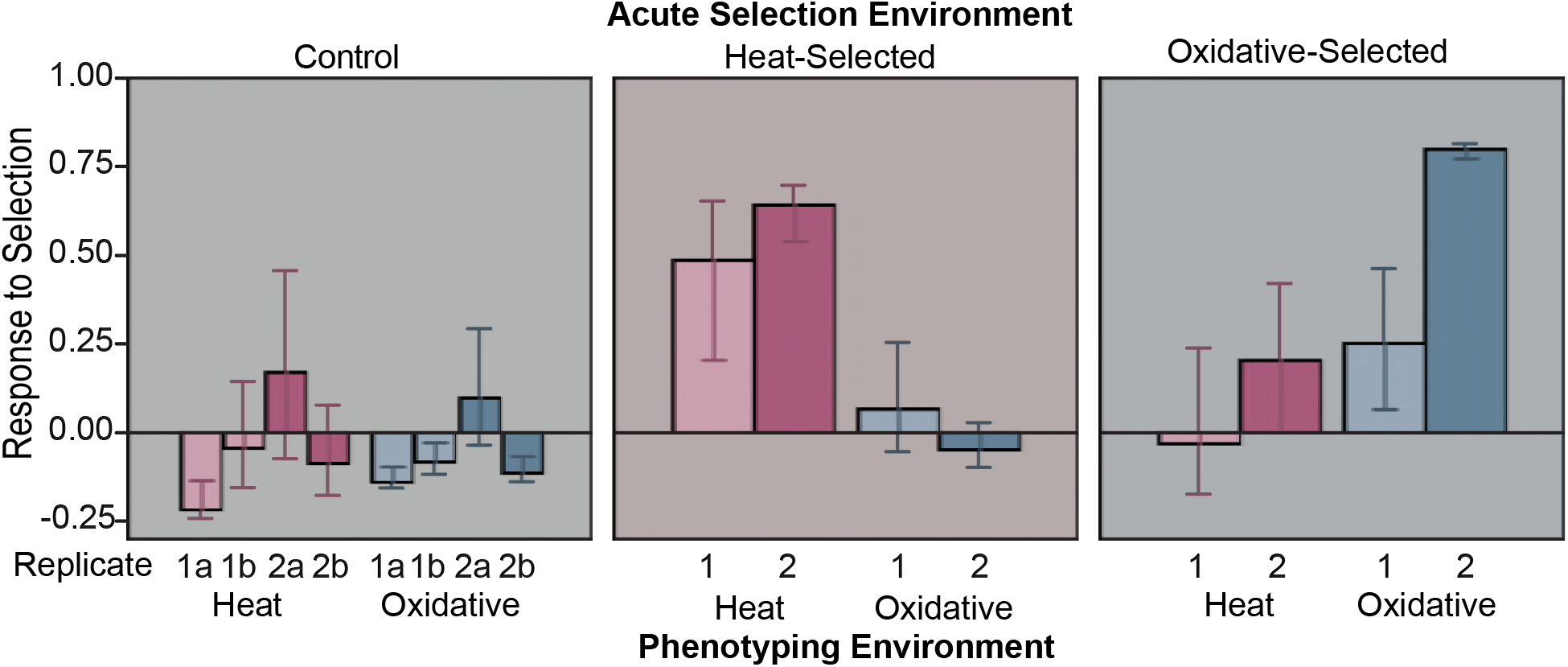
Direct and correlated response to selection for each replicate of experimentally evolved lines. A direct response to selection is measured by the response to selection where the phenotyping environment matches the acute selection environment and a correlated response to selection is measured by the response to selection where the phenotyping environment does not match the acute selection environment. There is a strong direct response to selection seen in both acute heat and oxidative selected populations but no correlated response to selection. Modified from Supplementary Figure S1 in (Sikkink *et al*. 2015).

## MATERIALS AND METHODS

### EXPERIMENTAL EVOLUTION

As described in (Sikkink, Reynolds, *et al*. 2014), the base “ancestral” population used to initiate the experimental evolution replicates was created using 26 isofemale strains, to create a population representative of the naturally segregating genetic variation of a single population of *C. remanei* collected in Ontario, Canada. This population (PX443) was used to establish each of the replicate selection and control lines and was frozen soon after creation to prevent adaptation to the lab environment. Using the methods described below, we determined that this population had high initial levels of genetic diversity (genome wide average π = 0.0178, Figure S1). All natural isolates and experimental lines were raised on Nematode Growth Medium-lite (NGM-lite, U.S. Biological) seeded with *Escherichia coli* strain OP50 (Brenner 1974).

Evolved populations of *C. remanei* belong to one of two replicate experimental blocks, each derived from an independently thawed subset of the ancestral population. Within each replicate block populations were evolved in parallel to one of three acute environments: control (2 lines/block), acute heat stress (1 line/block) and acute oxidative stress (1 line/block) as previously described (Sikkink, Reynolds, *et al*. 2014; Sikkink *et al*. 2015). Here, an acute stress is one that the worms experienced for four hours as L1 juveniles before returning to the laboratory control environment. The acute heat stress assay induced ~70% mortality in the ancestor population and the acute oxidative stress assay induced ~80% mortality in the ancestor population (Sikkink *et al*. 2015). Populations were subjected to acute stress conditions every two generations or whenever the population produced ≥ 24,000 eggs, whichever happened later. Control populations were randomly culled to 1000 worms during selective generations. For all acute stress events, each population (control evolved, acute heat stress evolved and acute oxidative stress evolved) was age synchronized via a bleach treatment (Stiernagle 2006). All experimentally evolved populations were subject to a total of ten acute stress events. As described in (Sikkink, Reynolds, *et al*. 2014) each population was frozen after about every second generation of acute stress selection to ensure that the worms did not lose the ability to survive a freeze and to provide a record of evolutionary changes. All told, there were a total of eight experimentally evolved populations created from the ancestor population: four control replicates, two acute heat stress replicates and two oxidative stress replicates.

### PREPARATION OF WHOLE GENOME SEQUENCING LIBRARIES

We determined the genetic response to selection using a population-based pooled sequencing (pool-seq) approach (Schlotterer *et al*. 2014). Ancestral and acute evolved populations were sequenced via whole genome shotgun sequencing. Approximately 10,000 L1 individuals (~24 hours old juvenile) were pooled together for DNA extraction and sequencing library preparation. Samples were prepared using the Nextera DNA Sample Preparation Kit (Illumina). Multiple sequencing libraries were prepared from the original DNA sample as needed to ensure adequate mean sequencing coverage (Table S1). Whole genome sequencing libraries were obtained for the ancestor and all eight evolved populations. Samples were sequenced on an Illumina HiSeq 2000 and Illumina HiSeq 2500 at the University of Oregon Genomics and Cell Characterization Core Facility. All sequences were deposited on NCBI’s Sequence Read Archive, accession number SRP126594 (https://www.ncbi.nlm.nih.gov/sra/SRP126594).

### WHOLE GENOME SEQUENCING ANALYSIS

Raw sequence reads were quality filtered using the ‘process_shortreads’ component of Stacks (Catchen *et al*. 2013). Low quality reads (mapping and base quality less than 20) were trimmed or discarded; all reads that passed quality filtering were aligned to the *C. remanei* reference genome assembled from the PX356 *C. remanei* inbred strain (Fierst *et al*. 2015) using the alignment program *GMAP – GSNAP* (Wu and Nacu 2010). Single nucleotide polymorphism (SNP) tolerant alignment parameters were used to ensure that divergent SNPs would align to the reference genome. We used available gene annotation data for PX356 (Fierst *et al*. 2015) in downstream data analysis. However, we also wanted to make use of the more extensive functional annotation data available for *C. elegans*. To that end, we used the software *OrthoFinder* (Emms and Kelly 2015) to identify *C. elegans* orthologs of *C. remanei* genes and protein domains. The *C. elegans* protein annotations were obtained from Wormbase ParaSite (Lee *et al*. 2018), version WS258.

We used the R (R Core Team 2017) package *Nest* (Jonas *et al*. 2016) to estimate effective population size (*N_e_*) of each experimental replicate using temporal changes in allele frequency based on existing theory (Krimbas and Tsakas 1971; Jorde and Ryman 2007; Waples *et al*. 2016), modified for samples taken from pooled population data. The input data, allele frequency and coverage data for each SNP at generation 0 (ancestor population) and generation 30 (each evolved population) was obtained from the program ‘snp-frequency-diff.pl’ from *Popoolation2* (Kofler, Pandey, *et al*. 2011). Only SNPs with a minimum coverage of 10X in each population were included in the effective population size estimator. Adding a maximum coverage threshold did not affect the estimated Ne. We used a pool sample size (the number of individuals who went into the original sequencing sample) of 1000 but increasing the pool size from 1000 to 10,000 individuals had a negligible effect on the estimated Ne. Expected Ne based on census population sizes for the duration of the experiment were calculated based on Equation 3.5 in (Kimura 1983) using the ‘harmonic.mean’ function in the R (R Core Team 2017) package *psych* (https://CRAN.R-project.org/package=psych).

Nucleotide diversity in the ancestor population was calculated over 1000 base pair (bp) windows using the program *Popoolation* (Kofler, Orozco-terWengel, *et al*. 2011). Allele frequency differences were analyzed on a site by site basis between the ancestor and control, heat stress evolved or oxidative stress evolved populations using the Fisher’s exact test program from the program *Popoolation2* (Kofler, Pandey, *et al*. 2011). Each SNP had to have a minimum of 10X coverage, a maximum coverage of 98% of total sequence coverage across the genome and the minor allele had to have at least two copies at a site in order to be retained for further analysis. Allele frequency differences were also analyzed over a range of sliding window sizes where each sliding window overlapped with one third of the length of the previous window. For each sliding window 75% or more of the bases in it had to meet the minimum and maximum coverage thresholds used for the single locus Fisher’s exact test comparison in order to be included. The geometric mean *p*-value of Fisher’s exact test across the window was calculated for each interval.

We then used a permutation test to determine whether or not a sliding window was significantly diverged from the ancestor population using a random sampling program in which windows were created at random by calculating the geometric mean *p*-value for a set of Fisher’s exact test outcomes from the pool of single-locus results (script available in File S1). The mean number of SNPs in an empirical sliding window of a particular size was used to determine the number of SNPs in each random sample window. Random sampling was repeated 1,600,000 times for each ancestor-experimentally evolved population pair. Random windows equivalent to empirical 1000–20,000 bp windows were created, with the 5000 bp window size chosen for the reported results because it is approximately the same size or slightly larger than nearly all genes in *C. remanei*, ensuring that genic and near genic regions could be captured in one window (see Figures S2 and S3 for a comparison of the distribution of empirical versus random sample windows). The random sampling program code can be found in File S1. Every ancestor-evolved population pair produced roughly 20,000 5 kp windows. We used the Bonferonni corrected *p*-value for 20,000 tests (2.5×10^−6^) as the cut-off for significance in each random sample window file. Any empirical sliding window that had a mean Fisher’s exact test value greater than or equal to the 99.99975% of the random sample window distribution was considered to be significantly differentiated from the ancestral population. A given window was defined as being significantly differentiated within a given selection regime (acute heat or oxidative stress selected) if it did not overlap with differentiated windows in the control populations and if the window was classified as significantly differentiated within both replicates. *BEDTools* ‘intersect’ (move Quinlan and Hall citation here) (Quinlan and Hall 2010) was used to identify overlapping genomic windows and genes under significantly differentiated genomic windows.

### HAPLOTYPE PHASING USING SINGLE WORM RAD SEQUENCING

We tested for the potential buildup of linkage disequilibrium within selection lines by crossing worms from the experimental evolution population with the reference genome strain (PX356), thereby generating F1s heterozygous for a single ancestral population haplotype. Single worm RAD sequencing libraries were prepared for 77 male F1 worms. Because the genotype of the reference strain is known, the haplotype of the target crossed individual could be directly determined in the F1 heterozygotes.

The RAD capture (Ali *et al*. 2016) method was used to genotype F1 worms. RAD capture is a flexible reduced-representation sequencing method that allowed us to multiplex and economically sequence all 77 genomes in a single Illumina lane. We digested genomic DNA with the restriction enzyme *EcoRI;* ligated digested fragments to barcoded, biotinylated adaptors; and sheared fragments with a BioRuptor. We enriched for fragments containing *EcoRI* cut sites with streptavidin beads and prepared libraries for sequencing with the GE GenomiPhi V3 DNA Amplification kit. Samples were sequenced on an Illumina HiSeq 4000 at the University of Oregon Genomics and Cell Characterization Core Facility.

Raw fastq files were reoriented using flip2BeRAD (https://github.com/tylerhether/Flip2BeRAD), which reorients read pairs such that all RAD tags and inline barcodes are present as read 1. Retained reads were filtered and demulitplexed with ‘process_radtags’ from *Stacks* using standard stringencies. Reads were aligned to the *C. remanei* PX356 reference sequence (Fierst *et al*. 2015) using *GSNAP* ultrafast settings. We only used read-pairs that mapped to the reference in the expected orientation (pairs facing inward toward each other) in downstream analyses. Individuals were genotyped using the ‘HaplotypeCaller’ and ‘GenotypeGVCF’ programs from *GATK* (https://software.broadinstitute.org/gatk/) (Van der Auwera *et al*. 2013). *VCFtools* (Danecek *et al*. 2011) was used to filter sites based on missing data, minor allele frequencies and genomic position. *PLINK* (http://pngu.mgh.harvard.edu/purcell/plink/; Purcell *et al*. 2007) was used to calculate linkage disequilibrium on the 13 largest contigs, which account for ~54% of total genome length. The squared inter-variant allele count correlations (r^2^ values) were reported from *PLINK* (Purcell *et al*. 2007).

### TRANSCRIPTIONAL PROFILING OF EVOLVED POPULATIONS

RNA sequencing libraries were prepared as described in (Sikkink, Reynolds, *et al*. 2014). Briefly, pooled samples of ~100,000 age-synchronized L1 larval worms were collected from the ancestor population, one of the control populations from the replicate two block (Control line – 2), and the acute heat stress and acute oxidative stress evolved populations from the replicate two block. Total RNA was isolated from each sample using standard TRIzol (Ambion) methods. All tissue samples were collected from L1 larval worms. All worms were raised in the standard lab environment of 20°C; the two stress evolved populations were not exposed to the stressor (either heat stress or oxidative stress) prior to tissue collection. We collected 6–8 replicate samples from each evolved line from at least two independently thawed populations from each evolved line, except for the oxidative stress evolved line, where 6 replicate samples were collected from one thaw. Samples were sequenced on an Illumina HiSeq 2000 at the University of Oregon Genomics and Cell Characterization Core Facility. All transcriptome sequences have been deposited in NCBI’s Gene Expression Omnibus (Edgar *et al*. 2001) and are accessible through GEO Series accession number GSE108235.

Raw sequence reads were quality filtered as described in (Sikkink, Ituarte, *et al*. 2014). Briefly, raw sequencing reads were quality filtered using the ‘process_shortreads’ component of *Stacks* (Catchen *et al*. 2013) and all reads that passed quality filtering were aligned to *a C. remanei* reference genome assembled from the PX356 *C. remanei* inbred strain (Fierst *et al*. 2015) using *GMAP – GSNAP* (Wu and Nacu 2010). We used the ‘htseq-count’ tool from the Python *HTSeq* package (Anders *et al*. 2014) to count all reads that aligned to protein–coding genes, using the criteria described in (Sikkink, Reynolds, *et al*. 2014).

Differential gene expression analysis was conducted using the R package *DESeq2* (Love *et al*. 2014). Low variance genes were filtered using *DESeq2*’s automatic filter. We tested for differences in gene expression between the ancestor and three experimentally evolved populations (control line – 2, acute heat stress evolved and acute oxidative stress evolved). Genes were only called as differentially expressed in each comparison if the *p*-value was less than 0.05 after Benjamini-Hochberg adjustment as implemented in *DESeq2*. We also tested for a relationship between log_2_ fold changes in gene expression between acute heat stress–ancestor and acute oxidative stress–ancestor comparisons using Model II linear regression. This analysis was implemented in the *lmodel2* package in R (R Core Team 2017).

### DATA AVAILABILITY

File S1 contains the python script used to create the random sampling windows. Files S2 and S3 contain the full list of genes identified as differentially expressed and/or divergent, including *C. elegans* ortholog names. Table S1 contains sequencing coverage for all experimental populations. Files S1-S3, Table S1 and all supplemental figures available on figshare (https://figshare.com/projects/Supplemental_material_for_Complex_pleiotropic_genetic_archite_cture_of_evolved_heat_stress_and_oxidative_stress_resistnace_in_the_nematode_Caenorhabditi_s_remanei_/58364). Whole genome sequencing data are available at NCBI SRA database under accession number PRJNA422140. The transcriptome data have been deposited in NCBI’s Gene Expression Omnibus (Edgar *et al*. 2001) and are accessible through GEO Series accession number GSE108235. This Whole Genome Shotgun project has been deposited at DDBJ/ENA/GenBank under the accession LFJK00000000. The version described in this paper is version LFJK02000000.

## RESULTS

### ADAPTATION TO ACUTE STRESS INVOLVES HUNDREDS OF LOCI

As demonstrated previously, selection for resistance to acute heat stress and acute oxidative stress leads to the evolution of significantly increased survival within the acute heat stress and acute oxidative stress environments (Sikkink *et al*. 2015). Using whole-genome resequencing of the selected lines, we find that this phenotypic response is strongly echoed across the entire genome: 13–27% of genomic windows within each population were more differentiated then expected by chance (Figure 2). There was a large amount of genetic differentiation between not only the ancestor and stress evolved populations, but also between the ancestor and control evolved populations (Figure 2). Because there was essentially no detectable response to either an acute heat stress or an acute oxidative stress in any of the control evolved populations at the phenotypic level (Figure 1), genetic differentiation within the control populations is most likely due to the effects of genetic drift and adaptation to the lab environment, since this population had very recently been brought into the lab from nature (Sikkink, Reynolds, *et al*. 2014). For this reason, we focused on differentiated genomic regions in the acute stress populations that were not differentiated in any of the control populations (that is, unique) and were shared among the evolutionary replicates (Figure 2). Those regions are referred to “differentiated windows” in what follows. After this filtering, we were able to identify hundreds of reproducibly differentiated windows: 495 and 491 differentiated 5 kp windows in the acute heat stress evolved and acute oxidative stress evolved populations, respectively.

**Figure 2.**
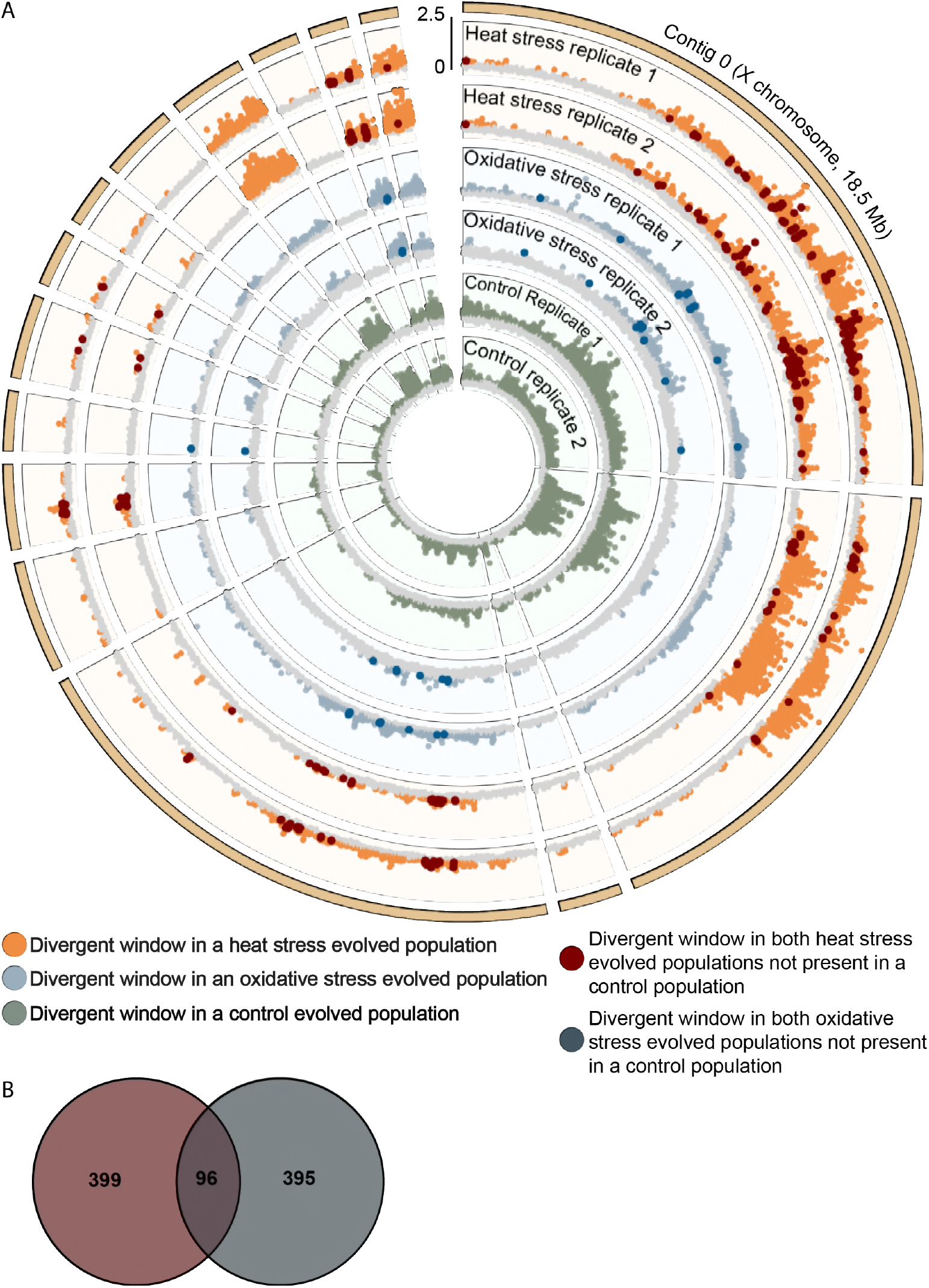
Location of significantly differentiated 5000 bp genomic windows. Mean −log10 of the Fisher’s Exact Test p-value for each window shown. The two control populations for each replicate are plotted on same ring. Widespread divergence from the ancestor (non-grey dots) is seen in all evolved populations, including the control populations. (b) Overlap between unique differentiated windows between heat stress and oxidative stress evolved populations. Only around one fifth of the unique differentiated windows overlap in location.

### GENOMIC DIFFERENTIATION IN HEAT STRESS AND OXIDATIVE STRESS EVOLVED POPULATIONS IS LARGELY INDEPENDENT

Stress resistance, including resistance to heat and oxidative stress, is expected have a common genetic basis in nematodes based on previous studies of *C. elegans* (Kenyon 2005; Rodriguez *et al*. 2013). In contrast to these expectations, we observed little correlated phenotypic response to either acute heat stress resistance or acute oxidative stress, even while there was a strong evolutionary response within a given selective environment (Figure 1; Sikkink *et al*. 2015). This lack of correlation at the phenotypic level is strongly reflected in the pattern of evolutionary response at the genetic level. We identified 491 differentiated windows in lines evolved under acute oxidative stress conditions and 495 in lines evolved under acute heat stress. Of these, only 96 are shared between the two selective regimes (Figure 2B). This indicates a general lack of pleiotropy in the genes underlying the response to selection for heat and oxidative stress.

### LINKAGE DISEQUILIBRIUM AS A POSSIBLE SOURCE OF COMPLEXITY IN GENETIC RESPONSE

The large number of genomic regions that we saw responding to selection could be due to a large number of genes underlying the traits of interest or, alternatively, due to the impact of linked selection on a smaller number of causative loci during adaptation (Franssen *et al*. 2015). We used single-individual SNP data in the ancestral population to estimate the degree of linkage disequilibrium within these populations. We found that linkage disequilibrium in the ancestor decays to background levels within 200 bp (Figures S4 and S5), very similar to the pattern observed within natural populations (Cutter *et al*. 2006), and found no evidence of local or long-distance linkage disequilibrium beyond that (Figures S6 and S7).

Even in the absence of haplotype structure in the ancestral population, selection itself may generate linked responses to selection within specific chromosomal regions, especially those with reduced recombination rates (Burri *et al*. 2015).

We cannot directly address this with pool-seq data, which is one limitation of our resequencing approach (Franssen *et al*. 2015). However, any form of linked selection would be expected to bias the results in favor of finding apparent pleiotropy, since linkage is another important source of genetic correlation among traits (Lande 1984). The larger the haplotype block, the higher the probability that two genes with independent effects on a suite of traits display a correlated response to selection. It is therefore likely that we have actually overestimated the degree of shared pleiotropy within the 5kb genomic windows of our analysis.

### EFFECTIVE POPULATION SIZE

Effective population size is an important parameter in experimental evolution, as effective population size is directly related to the relative opportunity and strength of random genetic drift versus selection in a population (Kimura 1983). Estimates of effective population size (*N_e_*) for each of the evolved populations ranged from 379-1602 (Table 1). This is much smaller than the expected *N_e_* given our census population sizes (Table 2), which fluctuated between 1,000–24,000 worms (Sikkink *et al*. 2015). We also obtained *N_e_* estimates using SNPs found only on the X chromosome. *C. remanei* has an XX/XO sex determination system (Hodgkin 1987; Thomas *et al*. 2012) so we would expect *N_e_* on the X chromosome to be around 75% of the *N_e_* on the autosomes. However, our estimates show X chromosome *N_e_* values that range from 60% to 146% of the all SNP *N_e_* estimates (Table 1).

**Table 1.** Estimated effective population (*N_e_*) size for each experimentally evolved population. *N_e_* estimated for all eight evolved populations using allele frequency and coverage data estimated using *Popoolation2* (Kofler, Pandey, *et al*. 2011). *N_e_* for all SNPs and just SNPs on the assembled X chromosome shown.

| Population | $N_e$ | X Chr $N_e$ |
| --- | --- | --- |
| Replicate 1 Control – 1 | 650 | 619 |
| Replicate 1 Control – 2 | 781 | 479 |
| Replicate 1 Heat stress | 978 | 640 |
| Replicate 1 Oxidative stress | 1602 | 1449 |
| Replicate 2 Control – 1 | 1051 | 1507 |
| Replicate 2 Control – 2 | 821 | 805 |
| Replicate 2 Heat stress | 654 | 528 |
| Replicate 2 Oxidative stress | 379 | 557 |

**Table 2.** Expected effective population sizes for the three experimental evolution conditions given the cyclical census population sizes described in Sikkink *et al*. (2015). Ranges come from different intermediate generation census population sizes. *N_e_* values calculated using equation 3.5 in Kimura (1983).

| Population | Expected $N_e$ |
| --- | --- |
| Control evolved | ~2,500 – ~2,800 |
| Heat stress evolved | ~5,000 – ~6100 |
| Oxidative stress evolved | ~3,500 – ~4600 |

### PLEIOTROPIC STRUCTURE OF GENE EXPRESSION

The lack of correlated response to selection in the stress resistance phenotypes is echoed at the level of the molecular phenotype of gene expression as well. In this case, however, many thousand responses can be interrogated simultaneously. Here, changes in expression are based on the comparison of baseline expression in the evolved populations versus those in the ancestor population. The numbers of differentially expressed genes greatly differ across selective environments (906 in the oxidative stress evolved population versus 91 in the heat stress evolved populations), and 21 genes are in common between the two groups (Figure 3). One of the common significantly differentially expressed genes is a putative member of the HSP70 family that promotes protein binding in response to stress and constitutively, but the other shared differentially expressed genes were not part of any canonical stress response pathways (File S2).

**Figure 3.**
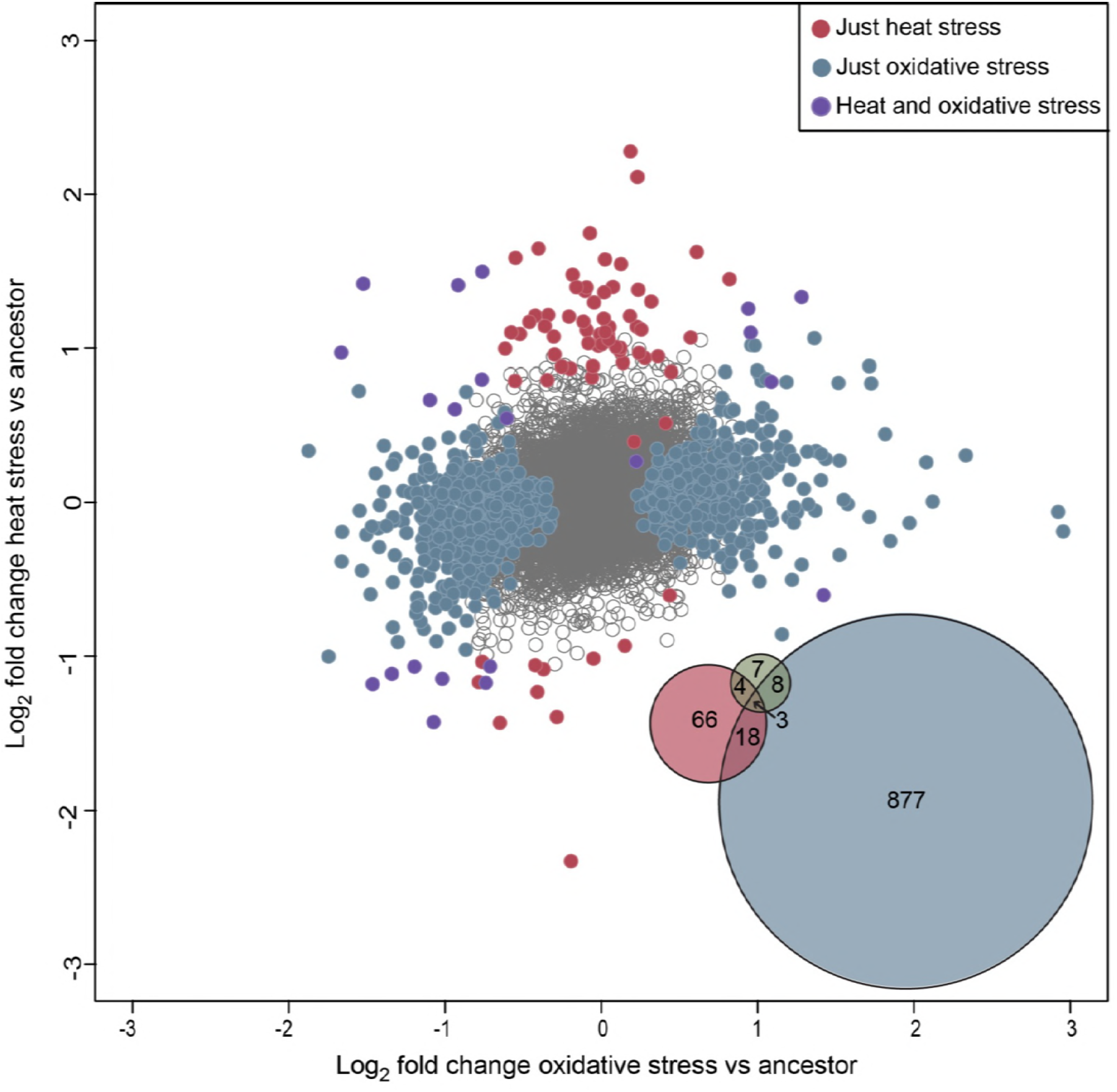
Correlation between log2 fold change of heat stress evolved vs. ancestor and oxidative stress evolved vs. ancestor tests for differential gene expression. The Venn diagram in the lower right shows the overlap in identity of significantly differentially expressed genes in the heat stress evolved, oxidative stress evolved and control evolved populations.

In addition, we observed an interesting pattern in the directionality of the evolution of gene expression. Genes differentially expressed in the heat stress evolved population were more likely to be up-regulated than down-regulated compared to both the oxidative stress evolved population (Fisher’s exact test, *p* = 1.7×10^−8^) and the control population (Fisher’s exact test, *p* = 1.9×10^−4^). However, the proportions of up-regulated versus down-regulated differentially expressed genes in the oxidative stress evolved population were not significantly different from the control population (Fisher’s exact test, *p* = 0.29). There was also a weak correlation between changes in gene expression between oxidative-stress evolved and heat-stress evolved populations (r^2^ = 0.049, *p* = 0.01), but this correlation appears to be largely driven by a relationship between the genes that have not changed in expression very much (i.e., it is a weak aggregate response over many genes).

### CANDIDATE GENES UNDER SELECTION FOR ACUTE HEAT STRESS AND ACUTE OXIDATIVE STRESS

Genes implicated as potentially being involved in the phenotypic response to selection, either via differences in gene expression or divergence due to allele frequency changes tended to create non-overlapping sets among the stress-response environments. Specifically, only three genes were found to be in common for the differential expression and genomic divergence gene sets within the heat stress evolved populations and 16 in common for the oxidative stress evolved populations. None of the shared genes had annotations associated with, or were orthologs of, genes known to be relevant to stress resistance (Files S2 and S3).

#### Differentiated genes due to allele frequency changes

There were 591 genes contained within significantly differentiated genomic windows in the heat stress evolved populations and 590 genes contained within significantly differentiated genomic regions in the oxidative stress evolved populations. Of those differentiated genes, 126 are shared in common between the heat stress and oxidative stress evolved populations. There are hundreds of other genes in these regions (File S3) and, again, for the most part they do not contain members of the canonical stress response pathways that have been determined via mutagenesis approaches.

We did identify a small number of stress response pathway candidate genes found within the differentiated genome regions in the heat stress evolved populations: a gene with a heat shock chaperonin-binding motif, *daf-8*, three insulin-like growth factor binding proteins, and *Ras* (File S3). The heat shock chaperonin-binding motif is found on the stress-inducible phosphoprotein STI1, known to bind heat stress proteins (Song *et al*. 2009). *daf-8* (Figure 4A) is a member of the TGF-β pathway, one of the signaling response pathways involved in heat shock response in *C. elegans* (Rodriguez *et al*. 2013). Insulin like growth factor binding proteins control the distribution and activity of insulin like growth factor receptor, an important stress response transmembrane receptor protein (Rodriguez *et al*. 2013; Murphy and Hu 2013). Finally, *Ras* is a member of the MAPK signaling pathway, which has been found to be responsive to pathogen (Ewbank 2006) and endoplasmic reticulum stress (Darling and Cook 2014). There is one candidate gene in the oxidative stress evolved populations with a known stress response function: SKI-interacting protein, which is a member of the TGF-β signaling pathway that promotes dauer formation when active (Savage-Dunn 2005).

**Figure 4.**
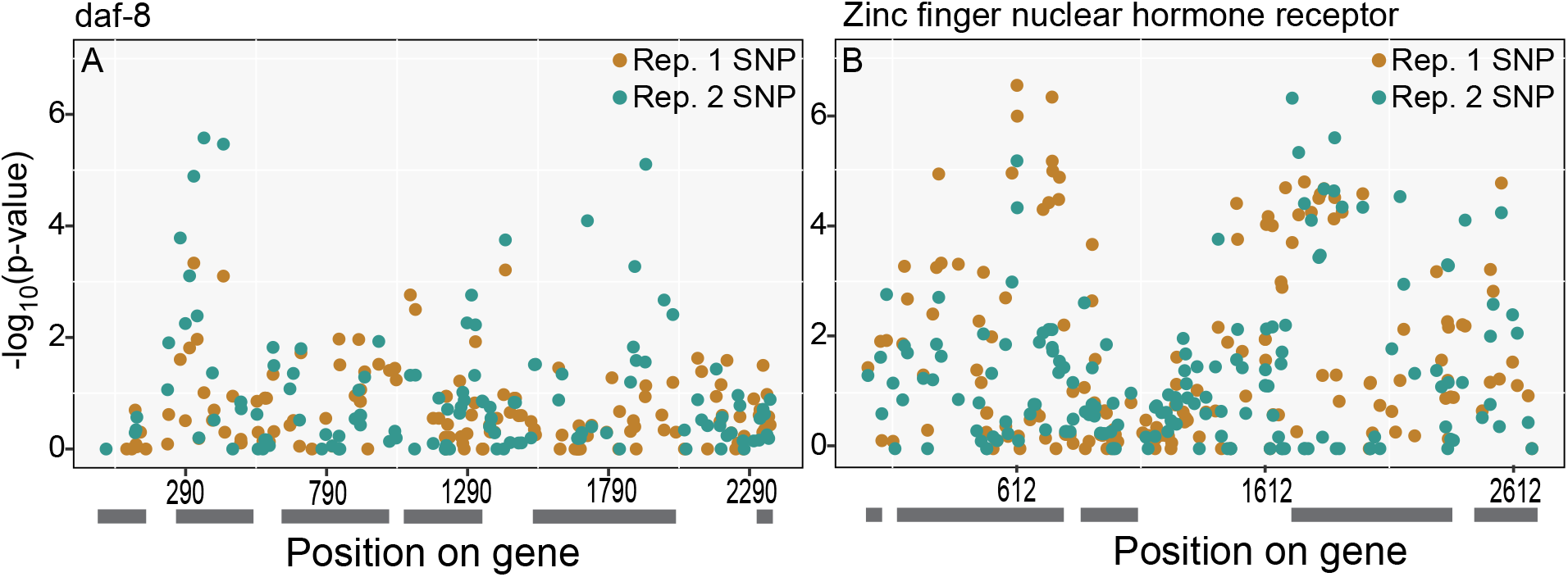
(a) −log10 *p*-values of Fisher’s Exact Test for each SNP in *daf-8* in the heat stress evolved populations. *daf-8* affects dauer development and is a component of the TGF-β signaling pathway (Savage-Dunn 2005). Grey bars indicate exons. (b) −log10 *p*-values of Fisher’ s Exact Test for each SNP in a zinc finger nuclear hormone receptor (*nhr-27*). This gene has lowest mean Fisher’s Exact Test *p*-value of the differentiated genes in the heat stress evolved populations. Grey bars indicate exons.

#### Differentially expressed genes

Similar to the pattern described above for differentiated genes, only a handful of the significantly differentially expressed genes have known or putative functions in abiotic stress response. There were significantly up-regulated heat shock proteins (HSPs) in both the heat shock evolved and oxidative shock evolved populations, and up-regulated insulin like growth factor binding proteins in the oxidative shock evolved population. Similar to previous work (Sikkink, Ituarte, *et al*. 2014), we find that up-regulated HSPs are significantly enriched in the heat stress evolved population (Fisher’s exact test, *p* = 0.00146), as well as in the oxidative stress evolved population (Fisher’s exact test, *p* = 0.029). In addition to up-regulated HSPs, SKN-1, a transcription factor whose regulatory targets are important for oxidative stress resistance (An and Blackwell 2003; Murphy and Hu 2013), was also significantly up-regulated in oxidative stress evolved population.

## DISCUSSION

Whole organism phenotypes are the combined result of the effects of and interactions among tens of thousands of genes. The classic assumption within the field of quantitative genetics is that majority of phenotypic variation found within natural populations is quantitative, with the phenotypic differences among individuals likely caused by a large number of genes operating within a complex environmental context (Hill 2009). Yet over the last two decades a small controversy has emerged about whether we expect the genetic basis of adaptation to be dominated by a few genes of major effect or whether it will be highly polygenic (Orr and Coyne 1992; Orr 2009; Rockman 2012). While this question is obviously most relevant in the context of evolution within specific natural populations, there is an unavoidable conflation of adaptation, demographic history, and chance events for any particular realization of an evolutionary lineage that makes it impossible to test specific hypotheses within a single instance. For example, even in the case of putatively adaptive armor loss in sticklebacks, in which it initially appeared that the same syndrome had been replicated many times independently during adaptation to fresh water, we now know that the genetic basis of this adaptation in each case is generated by a complex set of alleles that have likely been forged by thousands of generations of migration-selection balance (Colosimo *et al*. 2005; Nelson and Cresko 2018). Experimental evolution provides a means of addressing these questions in a controlled, repeatable manner. Our previous work has demonstrated that the nematode *C. remanei* is a powerful system for experimental evolution of stress resistance phenotypes (Sikkink, Reynolds, *et al*. 2014; Sikkink *et al*. 2015). Here, we use an evolve and re-sequence approach to probe the genetic basis of these phenotypic changes.

Our data demonstrate that both heat stress response and oxidative stress response are complex, polygenic traits, where hundreds of regions in the genome respond to selection (Figure 2) and hundreds of genes show significant changes in gene expression (Figure 3). This fits the pattern shown by many other experimental evolution and artificial selection studies (Turner *et al*. 2011; Orozco-terWengel *et al*. 2012; Pettersson *et al*. 2013; Hirsch *et al*. 2014), in which hundreds of differentiated genes and/or genomic regions have been identified that influence traits as varied as seed size, novel temperature regimes and body size.

The infinitesimal model, which allows researchers to predict the effects of selection on a trait without knowing the identity of the genes that underlie variation in that trait (Fisher 1918; Hill 2009), forms the cornerstone of quantitative genetics (Falconer and Mackay 1996) and has been very successful in plant and animal breeding (Hill 2014). Recent work in human genetics has found that that the vast majority of human traits and diseases are caused by variation in hundreds if not thousands of loci (Wood *et al*. 2014; Visscher *et al*. 2017). Taken in the context of this larger body of work, our study supports the view that much variation is quantitative in nature and responses to selection are generated by changes in hundreds of genes, even when selection itself is extremely strong (Johansson *et al*. 2010; Pettersson *et al*. 2013; Hirsch *et al*. 2014).

### THE GENOMIC BASIS OF MUTATIONAL PLEIOTROPY

Despite the expectation set by many molecular biology studies of stress response pathways in *C. elegans* (Kenyon 2010; Rodriguez *et al*. 2013; Murphy and Hu 2013), we found little evidence overall for pleiotropy for the genetic basis of acute heat stress resistance and acute oxidative stress resistance. There little to no correlated response to selection for heat stress or oxidative stress for the other stress at the level of survival (Figure 1 combine parentheses) (Sikkink *et al*. 2015) and mixed evidence for a correlated response for changes in gene expression (Figure 3). Finally, at the level of genomic response to selection, we also find little evidence for pleiotropy between the heat stress evolved and oxidative stress evolved populations at the level of individual genes (Figure 2).

This of course does not mean that pleiotropy is not a characteristic of any of the genes involved in stress response network, simply that it is not for those that respond to selection here. This discrepancy could be due to a number of different factors. First, a correlated response to selection is generated by the alleles that actually respond to selection, not all possible alleles for a gene (Lande 1979). It is possible that many of the stress response genes that have been identified via mutagenesis have broadly pleiotropic effects such that they cannot respond to direct selection for increased stress resistance per se because of potential negative pleiotropic effects on other traits (e.g., developmental rate, reproduction). That is, the genes that respond to selection are not major genes with many pleiotropic effects, since they would likely have some other negative pleiotropic effects. Second, molecular geneticists and quantitative geneticists focus on related but subtlety different definitions of the term pleiotropy. Most fundamentally, pleiotropy is a property of alleles, not genes (Phillips and McGuigan 2006), and so it is possible for there to be variation in pleiotropy at a single locus such that some segregating alleles will lead to a correlated response to selection while other will not. For stress resistance, we do not find a clear signal for canonical stress response pathway members in the first place, and so variance in pleiotropy at genes defined via measures of molecular pleiotropy still does not appear to be the primary driver here, regardless of any potential variation in allelic effects.

### THE GENETIC AND GENOMIC BASIS OF THE RESPONSE TO SELECTION

While there are a handful of genetically differentiated and differentially expressed genes with functions that are directly related to abiotic stress response, it is clear that the response to acute heat stress and acute oxidative stress cannot be explained by these genes alone. We do not find evidence that many of the canonical stress response genes, including *hsf-1* or *daf-16* (Rodriguez *et al*. 2013; Murphy and Hu 2013) for heat stress and *pink-1, lrk-1* and *sod-1, −2* or *−3* for oxidative stress (Rodriguez *et al*. 2013) responded to selection for acute heat stress or acute oxidative stress survival, although *skn-1* is constitutively up-regulated in the oxidative stress evolved population. This is further evidence that gene knockouts can paint an incomplete picture of the genetic architecture of a phenotype of interest and the fact that the genes that respond to selective pressures in a population are unlikely to be highly pleiotropic genes of large effect.

We found hundreds of genomic regions that appeared to respond to selection for either acute heat stress or acute oxidative stress resistance. If we assume that regions are independent of each other (i.e. in linkage equilibrium) then our results support the quantitative genetics worldview that variation in most phenotypes is due to many alleles, presumably of small effect (Falconer and Mackay 1996). However, some clustering of the genomic response can be clearly seen in Figure 2. For example, 22% of the differentiated windows in the two heat stress evolved populations are found on the X chromosome. When we look at strongly selected genes we do see evidence of local selective sweeps, where multiple SNPs have similarly highly significant Fisher’s exact test *p*-values and changes in allele frequency (Figure 4B, Figure S8). This raises the possibility that linkage disequilibrium is responsible for some of the response to selection in the evolved populations. Although we found that linkage disequilibrium decays to background levels within 200 bp within the ancestor population (Figures S4 and S5), it is possible that linkage disequilibrium developed over the course of the experimental evolution project, due to small effective population sizes (Table 1). However, the same clustering of genomic response is not seen in either of the oxidative stress evolved populations (Figure 2), where at most 10% of differentiated windows were found on one contig (that contig not shown in Figure 2).

We find that most our estimated effective population sizes are about an order of magnitude smaller than the post-selection census population size (Tables 1 and 2 combine parentheses) (Sikkink *et al*. 2015). Fluctuating population sizes will reduce *N_e_* (Kimura 1983), and in our case very strong selection has the potential to strongly reduce the level of variation at linked sites via the Hill-Robertson effect (Hill *et al*. 1966; Charlesworth 1996; Comeron *et al*. 2007; Charlesworth 2012). Even so, the reduction in Ne seems to be particularly severe relative to the very large population sizes (for an animal) carefully maintained throughout the experiment. For example, despite maintaining a census population size 10-20X larger than similar experiments in *Drosophila melanogaster*, our estimated *N_e_* values are only two–three times higher than estimated *N_e_* values for those populations (Orozco-terWengel *et al*. 2012; Jonas *et al*. 2016). Differences in overall strength of selection may play a role here (strength of selection is unknown in Jonas *et al*. 2016), but one additional source of reduction in Ne is potential variance in mating such that male mating success and/or female reproductive output displays a greater than Poisson variance (Wright 1931; Kimura 1983). This seems particularly likely because the control populations have similar effective population sizes to the selected lines. Variation in male mating ability has been previously documented within *C. remanei* (Palopoli *et al*. 2015) and bears further investigation within the context of experimental evolution within this species.

## Conclusion

Organisms are complex, integrated systems and the manner in which natural variation percolates though that system is likely to be highly structured by functional interactions. Although pleiotropy may be ubiquitous throughout the genome, individuals with mutations in highly pleiotropic genes are likely to suffer from reduced fitness since those mutations will most likely affect more traits than those under selection. As such, seeking to understand the genetic architecture of traits solely via an analysis of gene knockouts is likely to leave an incomplete picture of the overall structure of the pleiotropic network, especially with respect to segregating variation within natural populations. Using standing genetic variation, experimental evolution and whole genome sequencing therefore provides a powerful toolkit for understanding the genetic architecture of complex traits, no matter how complex they ultimately prove to be.

## Acknowledgments

We thank Sally E. Claridge and Erik Johnson for their help setting up crosses and samples for single worm RAD sequencing. We thank Gavin Woodruff for creating a script to run OrthoFinder. We also thank John Willis for creating the single worm RADseq libraries. This work was supported grants from the National Science Foundation (DEB-1210922 to WAC and KLS.; DEB-1607194 to PCP and CHO), grants from the National Institutes of Health (GM096008 and GM102511 to P.C.P.; RR032670 to W.A.C.), and pre-doctoral fellowships to CHO from the National Science Foundation and National Institutes of Health (T32 GM007413).

## Literature Cited

Ali, O. A., S. M. O’Rourke, S. J. Amish, M. H. Meek, G. Luikart et al., 2016 RAD Capture (Rapture): Flexible and Efficient Sequence-Based Genotyping. Genetics 202: 389–400.

An, J. H., and T. K. Blackwell, 2003 SKN-1 links *C. elegans* mesendodermal specification to a conserved oxidative stress response. Genes & Development 17: 1882–1893.

Anders, S., P. T. Pyl, and W. Huber, 2014 HTSeq—a Python framework to work with high-throughput sequencing data. Bioinformatics 31: btu638–169.

Arnold, S. J., 1992 Constraints on phenotypic evolution. Am Nat 140 Suppl 1: S85–107.

Arnold, S. J., R. Bürger, P. A. Hohenlohe, B. C. Ajie, and A. G. Jones, 2008 UNDERSTANDING THE EVOLUTION AND STABILITY OF THE G-MATRIX. Evolution 62: 2451–2461.

Barton, N. H., and M. Turelli, 1989 Evolutionary quantitative genetics: how little do we know? Annu. Rev. Genet. 23: 337–370.

Boyle, E. A., Y. I. Li, and J. K. Pritchard, 2017 An Expanded View of Complex Traits: From Polygenic to Omnigenic. Cell 169: 1177–1186.

Brenner, S., 1974 The genetics of *Caenorhabditis elegans*. Genetics 77: 71–94.

Burri, R., A. Nater, T. Kawakami, C. F. Mugal, P. I. Olason et al., 2015 Linked selection and recombination rate variation drive the evolution of the genomic landscape of differentiation across the speciation continuum of *Ficedula* flycatchers. Genome Research 25: 1656–1665.

Carroll, S. B., 2008 Evo-Devo and an Expanding Evolutionary Synthesis: A Genetic Theory of Morphological Evolution. Cell 134: 25–36.

Castillo, D. M., M. K. Burger, C. M. Lively, and L. F. Delph, 2015 Experimental evolution: Assortative mating and sexual selection, independent of local adaptation, lead to reproductive isolation in the nematode Caenorhabditis remanei. Evolution 69: 3141–3155.

Catchen, J., P. A. Hohenlohe, S. Bassham, A. Amores, and W. A. Cresko, 2013 Stacks: an analysis tool set for population genomics. Molecular Ecology 22: 3124–3140.

Charlesworth, B., 1996 Background selection and patterns of genetic diversity in Drosophila melanogaster. Genet. Res. 68: 131–149.

Charlesworth, B., 1990 Optimization Models, Quantitative Genetics, and Mutation. Evolution 44: 520–538.

Charlesworth, B., 2012 The Effects of Deleterious Mutations on Evolution at Linked Sites. Genetics 190: 5–22.

Cheverud, J. M., 1984 Quantitative genetics and developmental constraints on evolution by selection. J. Theor. Biol. 110: 155–171.

Colosimo, P. F., K. E. Hosemann, S. Balabhadra, G. Villarreal Jr, M. Dickson et al., 2005 Widespread Parallel Evolution in Sticklebacks by Repeated Fixation of Ectodysplasin Alleles. Science 307: 1928–1933.

Comeron, J. M., A. Williford, and R. M. Kliman, 2007 The Hill–Robertson effect: evolutionary consequences of weak selection and linkage in finite populations. Heredity 100: 19–31.

Consortium, S. W. G. O. T. P. G., S. Ripke, B. M. Neale, A. Corvin, J. T. R. Walters et al., 2014 Biological insights from 108 schizophrenia-associated genetic loci. Nature 511: 421–17.

Cutter, A. D., S. E. Baird, and D. Charlesworth, 2006 High Nucleotide Polymorphism and Rapid Decay of Linkage Disequilibrium in Wild Populations of *Caenorhabditis remanei*. Genetics 174: 901–913.

Danecek, P., A. Auton, G. Abecasis, C. A. Albers, E. Banks et al., 2011 The variant call format and VCFtools. Bioinformatics 27: 2156–2158.

Darling, N. J., and S. J. Cook, 2014 The role of MAPK signalling pathways in the response to endoplasmic reticulum stress. BBA - Molecular Cell Research 1843: 2150–2163.

Edgar, R., M. Domrachev, and A. E. Lash, 2001 Gene Expression Omnibus: NCBI gene expression and hybridization array data repository. Nucleic Acids Res. 30: 207–210.

Emms, D. M., and S. Kelly, 2015 OrthoFinder: solving fundamental biases in whole genome comparisons dramatically improves orthogroup inference accuracy. Genome Biol. 16: 157.

Ewbank, J., 2006 Signaling in the immune response. WormBook 1–12.

Falconer, D. S., and T. F. C. Mackay, 1996 Introduction to Quantitative Genetics. Harlow.

Fierst, J. L., J. H. Willis, C. G. Thomas, W. Wang, R. M. Reynolds et al., 2015 Reproductive Mode and the Evolution of Genome Size and Structure in *Caenorhabditis* Nematodes. (M. Blaxter, Ed.). PLoS Genetics 11: e1005323.

Fisher, R. A., 1918 The Correlation between Relative on the Supposition of Mendelian Inheritance. Transactions of the Royal Society of Edinburgh 52: 399–433.

Franssen, S. U., V. Nolte, R. Tobler, and C. Schlötterer, 2015 Patterns of linkage disequilibrium and long range hitchhiking in evolving experimental Drosophila melanogaster populations. Molecular Biology and Evolution 32: 495–509.

Hill, W. G., 2014 Applications of Population Genetics to Animal Breeding, From Wright, Fisher and Lush to Genomic Prediction. Genetics 196: 1–16.

Hill, W. G., 2009 Understanding and using quantitative genetic variation. Philos. Trans. R. Soc. Lond., B, Biol. Sci. 365: 73–85.

Hill, W. G., Robertson, A., and A. Robertson, 1966 The effect of linkage on limits to artificial selection. Genet. Res. 8: 269–294.

Hirsch, C. N., S. A. Flint-Garcia, T. M. Beissinger, S. R. Eichten, S. Deshpande et al., 2014 Insights into the Effects of Long-Term Artificial Selection on Seed Size in Maize. Genetics 198: 409–421.

Hodgkin, J., 1987 Primary sex determination in the nematode. Development, Suppl. 5–15.

Johansson, A. M., M. E. Pettersson, P. B. Siegel, and Ö. Carlborg, 2010 Genome-Wide Effects of Long-Term Divergent Selection (B. Walsh, Ed.). PLoS Genetics 6: e1001188.

Jonas, A., T. Taus, C. Kosiol, C. Schlötterer, and A. Futschik, 2016 Estimating the Effective Population Size from Temporal Allele Frequency Changes in Experimental Evolution. Genetics 204: 723–735.

Jones, A. G., R. B. U. rger, and S. J. Arnold, 2014 Epistasis and natural selection shape the mutational architecture of complex traits. Nature Communications 5: 1–10.

Jorde, P. E., and N. Ryman, 2007 Unbiased Estimator for Genetic Drift and Effective Population Size. Genetics 177: 927–935.

Jovelin, R., B. C. Ajie, and P. C. Phillips, 2003 Molecular evolution and quantitative variation for chemosensory behaviour in the nematode genus *Caenorhabditis*. Molecular Ecology 12: 1325–1337.

Jovelin, R., J. S. Comstock, A. D. Cutter, and P. C. Phillips, 2014 A recent global selective sweep on the age-1 phosphatidylinositol 3-OH kinase regulator of the insulin-like signaling pathway within Caenorhabditis remanei. G3&#58; Genes|Genomes|Genetics 4: 1123–1133.

Kemper, K. E., S. J. Saxton, S. Bolormaa, B. J. Hayes, and M. E. Goddard, 2014 Selection for complex traits leaves little or no classic signatures of selection. BMC Genomics 15: 1–14.

Kenyon, C., 2005 The Plasticity of Aging: Insights from Long-Lived Mutants. Cell 120: 449–460.

Kenyon, C. J., 2010 The genetics of ageing. Nature 464: 504–512.

Kimura, M., 1983 The neutral theory of molecular evolution. New York City.

Knight, C. G., N. Zitzmann, S. Prabhakar, R. Antrobus, R. Dwek et al., 2006 Unraveling adaptive evolution: how a single point mutation affects the protein coregulation network. Nat Genet 38: 1015–1022.

Kofler, R., and C. Schlötterer, 2013 A guide for the design of evolve and resequencing studies. Molecular Biology and Evolution 31: 474–483.

Kofler, R., P. Orozco-terWengel, N. De Maio, R. V. Pandey, V. Nolte et al., 2011 PoPoolation: a toolbox for population genetic analysis of next generation sequencing data from pooled individuals. (M. Kayser, Ed.). PLoS ONE 6: e15925.

Kofler, R., R. V. Pandey, and C. Schlötterer, 2011 PoPoolation2: identifying differentiation between populations using sequencing of pooled DNA samples (Pool-Seq). Bioinformatics 27: 3435–3436.

Krimbas, C. B., and S. Tsakas, 1971 The Genetics of Dacus Oleae. v. Changes of Esterase Polymorphism in a Natural Population Following Insecticide Control-Selection or Drift. Evolution 25: 454–460.

Lande, R., 1979 Quantitative genetic analysis of multivariate evolution, applied to brain: body size allometry. Evolution 33: 402–416.

Lande, R., 1984 The genetic correlation between characters maintained by selection, linkage and inbreeding. Genetical Research 44: 309–320.

Lango Allen, H., K. Estrada, G. Lettre, S. I. Berndt, M. N. Weedon et al., 2010 Hundreds of variants clustered in genomic loci and biological pathways affect human height. Nature 467: 832–838.

Lee, R. Y. N., K. L. Howe, T. W. Harris, V. Arnaboldi, S. Cain et al., 2018 WormBase 2017: molting into a new stage. Nucleic Acids Res. 46: D869–D874.

Linnen, C. R., Y.-P. Poh, B. K. Peterson, R. D. H. Barrett, J. G. Larson et al., 2013 Adaptive evolution of multiple traits through multiple mutations at a single gene. Science 339: 1312–1316.

Love, M. I., W. Huber, and S. Anders, 2014 Moderated estimation of fold change and dispersion for RNA-seq data with DESeq2. Genome Biol. 15:.

Murphy, C. T., and P. T. Hu, 2013 Insulin/insulin-like growth factor signaling in *C. elegans*. WormBook 1–43.

Natarajan, C., N. Inoguchi, R. E. Weber, A. Fago, H. Moriyama et al., 2013 Epistasis Among Adaptive Mutations in Deer Mouse Hemoglobin. Science 340: 1324–1327.

Nelson, T. C., and W. A. Cresko, 2018 Ancient genomic variation underlies repeated ecological adaptation in young stickleback populations. Evolution Letters 2: 9–21.

Nolte, I. M., P. J. van der Most, B. Z. Alizadeh, P. I. de Bakker, H. M. Boezen et al., 2017 Missing heritability: is the gap closing? An analysis of 32 complex traits in the Lifelines Cohort Study. Eur. J. Hum. Genet. 25: 877–885.

Orozco-terWengel, P., M. Kapun, V. Nolte, R. Kofler, T. Flatt et al., 2012 Adaptation of Drosophila to a novel laboratory environment reveals temporally heterogeneous trajectories of selected alleles. Molecular Ecology 21: 4931–4941.

Orr, H. A., 2009 Fitness and its role in evolutionary genetics. Nat. Rev. Genet. 10: 531–539.

Orr, H. A., and J. A. Coyne, 1992 The genetics of adaptation: a reassessment. Am Nat 140: 725–742.

Paaby, A. B., and M. V. Rockman, 2013 The many faces of pleiotropy. Trends in Genetics 29: 66–73.

Palopoli, M. F., C. Peden, C. Woo, K. Akiha, M. Ary et al., 2015 Natural and experimental evolution of sexual conflict within *Caenorhabditis* nematodes. BMC Evolutionary Biology 1–13.

Pettersson, M. E., A. M. Johansson, P. B. Siegel, and Ö. Carlborg, 2013 Dynamics of adaptive alleles in divergently selected body weight lines of chickens. G Genes Genomics Genetics 3: 2305–2312.

Phillips, P. C., 2005 Testing hypotheses regarding the genetics of adaptation. Genetica 123: 15–24.

Phillips, P. C., and K. L. McGuigan, 2006 Evolution of Genetic Variance-Covariance Structure, pp. 310–325 in Evolutionary Genetics, edited by C. W. Fox and J. B. Wolf.

Purcell, S., B. Neale, K. Todd-Brown, L. Thomas, M. A. R. Ferreira et al., 2007 PLINK: A Tool Set for Whole-Genome Association and Population-Based Linkage Analyses. The American Journal of Human Genetics 81: 559–575.

Quinlan, A. R., and I. M. Hall, 2010 BEDTools: a flexible suite of utilities for comparing genomic features. Bioinformatics 26: 841–842.

R Core Team, R. C., 2017 R: A language and environment for statistical computing.

Reynolds, R. M., and P. C. Phillips, 2013 Natural variation for lifespan and stress response in the nematode *Caenorhabditis remanei*. (M. Kaeberlein, Ed.). PLoS ONE 8: e58212.

Rockman, M. V., 2012 The QTN program and the alleles that matter for evolution: all that’s gold does not glitter. Evolution 66: 1–17.

Rodriguez, M., L. B. Snoek, M. De Bono, and J. E. Kammenga, 2013 Worms under stress: *C. elegans* stress response and its relevance to complex human disease and aging. Trends in Genetics 29: 367–374.

Savage-Dunn, C., 2005 TGF-β signaling. WormBook 1–12.

Schlotterer, C., R. Kofler, E. Versace, R. Tobler, C. Schlötterer et al., 2015 Combining experimental evolution with next-generation sequencing: a powerful tool to study adaptation from standing genetic variation. Heredity 114: 431–440.

Schlotterer, C., Schlötterer, C., R. Tobler, R. Kofler, and V. Nolte, 2014 Sequencing pools of individuals — mining genome-wide polymorphism data without big funding. Nat. Rev. Genet. 15: 749–763.

Sikkink, K. L., C. M. Ituarte, R. M. Reynolds, W. A. Cresko, and P. C. Phillips, 2014 The transgenerational effects of heat stress in the nematode *Caenorhabditis remanei* are negative and rapidly eliminated under direct selection for increased stress resistance in larvae. Genomics 104: 438–446.

Sikkink, K. L., R. M. Reynolds, W. A. Cresko, and P. C. Phillips, 2015 Environmentally induced changes in correlated responses to selection reveal variable pleiotropy across a complex genetic network. Evolution 69: 1128–1142.

Sikkink, K. L., R. M. Reynolds, C. M. Ituarte, W. A. Cresko, and P. C. Phillips, 2014 Rapid evolution of phenotypic plasticity and shifting thresholds of genetic assimilation in the nematode *Caenorhabditis remanei*. G Genes Genomics Genetics 4: 1103–1112.

Song, H.-O., W. Lee, K. An, H.-S. Lee, J. H. Cho et al., 2009 *C. elegans* STI-1, the Homolog of Sti1/Hop, Is Involved in Aging and Stress Response. Journal of Molecular Biology 390: 604–617.

Steiner, C. C., J. N. Weber, and H. E. Hoekstra, 2007 Adaptive variation in beach mice produced by two interacting pigmentation genes. (M. A. F. Noor & M. A. F. Noor, Eds.). PLoS Biol 5: e219.

Stiernagle, T., 2006 Maintenance of C. elegans. WormBook 1–11.

Teotónio, H., S. Estes, P. C. Phillips, and C. F. Baer, 2017 Experimental Evolution with *Caenorhabditis* Nematodes. Genetics 206: 691–716.

Thomas, C. G., G. C. Woodruff, and E. S. Haag, 2012 Causes and consequences of the evolution of reproductive mode in *Caenorhabditis* nematodes. Trends Genet. 28: 213–220.

Turner, T. L., A. D. Stewart, A. T. Fields, W. R. Rice, and A. M. Tarone, 2011 Population-based resequencing of experimentally evolved populations reveals the genetic basis of body size variation in Drosophila melanogaster. (G. Gibson, Ed.). PLoS Genetics 7: e1001336.

Van der Auwera, G. A., M. O. Carneiro, C. Hartl, R. Poplin, G. Del Angel et al., 2013 From FastQ data to high confidence variant calls: the Genome Analysis Toolkit best practices pipeline. Curr Protoc Bioinformatics 43: 11.10.1–33.

Visscher, P. M., N. R. Wray, Q. Zhang, P. Sklar, M. I. McCarthy et al., 2017 10 Years of GWAS Discovery: Biology, Function, and Translation. Am. J. Hum. Genet. 101: 5–22.

Wagner, G. P., and J. Zhang, 2011 The pleiotropic structure of the genotype–phenotype map: the evolvability of complex organisms. Nature Publishing Group 12: 204–213.

Waples, R. K., W. A. Larson, and R. S. Waples, 2016 Estimating contemporary effective population size in non-model species using linkage disequilibrium across thousands of loci. Heredity 117: 1–8.

Wood, A. R., T. Esko, J. Yang, S. Vedantam, T. H. Pers et al., 2014 Defining the role of common variation in the genomic and biological architecture of adult human height. Nat Genet 46: 1173–1186.

Wright, S., 1968 Evolution and the Genetics of Populations Volume I: Genetic and Biometric Foundations. Chicago.

Wright, S., 1931 Evolution in medelian populations. Genetics 16: 97–159.

Wu, T. D., and S. Nacu, 2010 Fast and SNP-tolerant detection of complex variants and splicing in short reads. Bioinformatics 26: 873–881.

